# Dynamic cytoplasmic fluidity during morphogenesis in a human fungal pathogen

**DOI:** 10.1101/2024.11.16.623909

**Authors:** A. Serrano, C. Puerner, E. Plumb, L. Chevalier, J. Elferich, L.R. Sinn, N. Grigorieff, M. Ralser, M. Delarue, M. Bassilana, R.A. Arkowitz

**Author notes:** To whom correspondence should be sent.

## Abstract

The molecular crowding of the cytoplasm impacts a range of cellular processes. Using a fluorescent microrheological probe (GEMs), we observed a striking decrease in molecular crowding during the yeast to filamentous growth transition in the human fungal pathogen *Candida albicans*. This decrease in crowding is due to a decrease in ribosome concentration that results in part from an inhibition of ribosome biogenesis, combined with an increase in cytoplasmic volume; leading to a dilution of the major cytoplasmic crowder. Moreover, our results suggest that inhibition of ribosome biogenesis is a trigger for *C. albicans* morphogenesis.

---

The cytoplasm is a crowded environment and molecular crowding can affect a range of biological functions^1,2^, including chemical reaction rates, protein complex formation and rates of cytoskeletal protein polymerization^3-5^. In the budding yeast *Saccharomyces cerevisiae* and mammalian cells the target of rapamycin complex (TORC) was shown to regulate ribosome concentration, and inhibition of TORC1 resulted in increased cytoplasmic fluidity^6^. More recently, cell cycle arrest mutants were shown to result in decreased macromolecular crowding of the cytoplasm, in part *via* ribosome downregulation^7^. Little is known about the relationship between molecular crowding in the cytoplasm and morphological growth states, for example when the human fungal pathogen *Candida albicans* switches from an ovoid yeast form to a filamentous hyphal form, a transition essential for virulence. Here, we show that there is a dramatic decrease in molecular crowding during filamentous growth, which is mediated by inhibition of ribosome biogenesis and subsequent dilution of this critical cytoplasmic crowder as a result of new growth. We propose that ribosome levels are tuned to regulate crowding in distinct growth states in this fungal pathogen. Furthermore, our results highlight that despite the new growth that occurs during the yeast-to hyphal transition, there is a substantial decrease in ribosome levels.

To investigate the link between cytoplasmic diffusivity at the mesoscale and cell morphology we took advantage of passive microrheological probes^6^, for which we can measure dynamics both in budding and hyphal cells. We used *C. albicans* cells expressing 40-nm GEMs. We imaged every 30 msec to obtain a signal sufficient for single particle tracking. The mean track length was 10 frames, which corresponds to 300 msec of imaging. The effective diffusion coefficient, D_eff_, which is inversely proportional to microviscosity for Brownian motion, was determined from the first 120 msec of acquisition. Temporal projections that were false colored for GEM distribution in budding cells, fixed budding cells and cells treated with fetal bovine serum for 60 min, in which a filament is evident, are shown in Figure 1A, with trajectories of GEMs in representative budding and filamentous cells. Fixation reduced the GEM D_eff_ of cells by over 10-fold. The median effective coefficient of diffusion in budding cells was ∼ 0.1 μm^2^ s^-1^ (Fig. 1B). This value was independent of GEM expression level, with similar D_eff_ observed with *TEF1* and *ADH1* promoter driven expression (Fig. S2), and somewhat lower than that observed in *S. cerevisiae*^6^.

**Figure 1.**
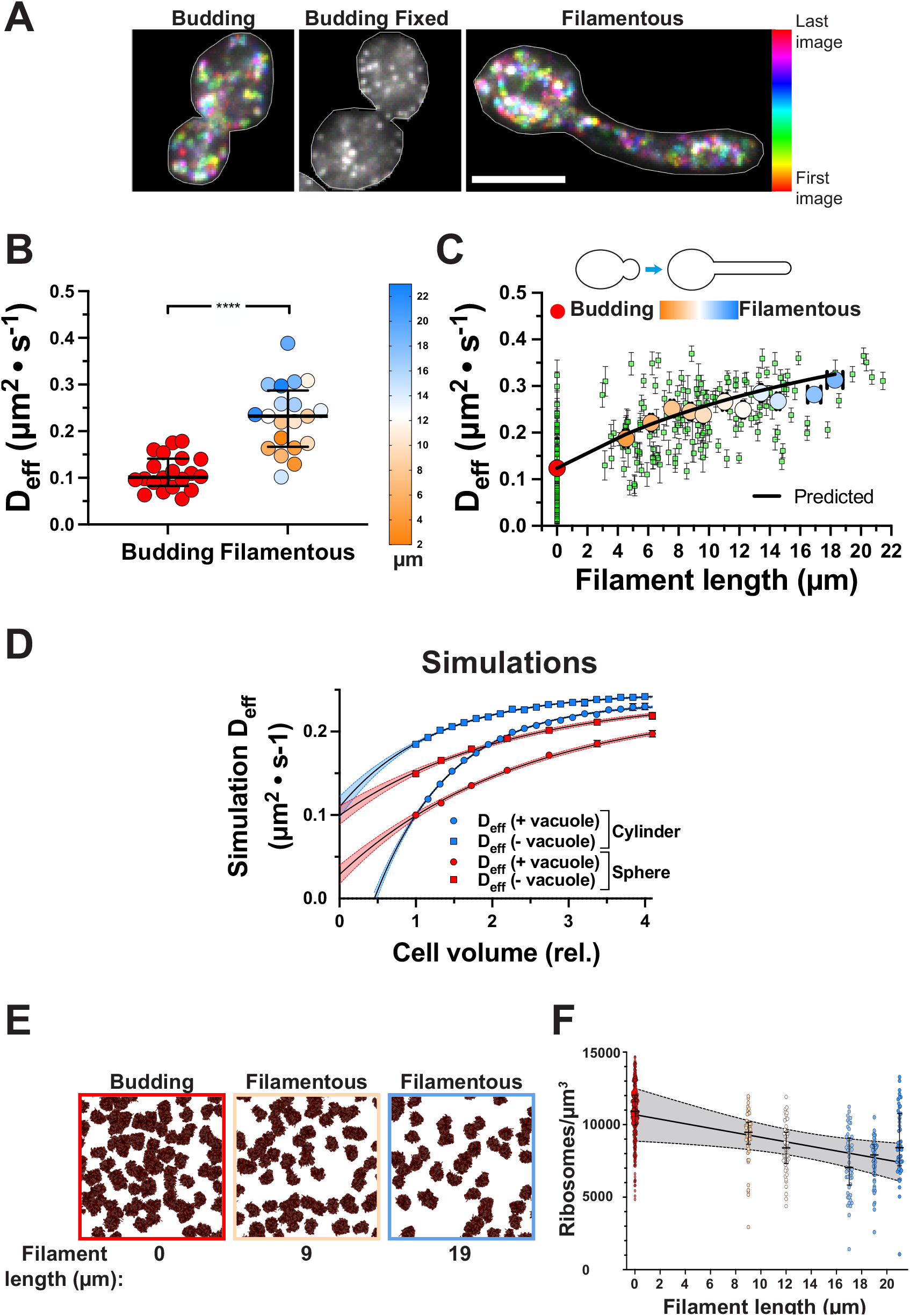
Cytoplasmic fluidity increases with filament length. A) Temporally color-coded projection of GEM dynamics. Indicated cells were imaged, 250 × 10 msec. B) Dramatic increase in GEM D_eff_ in filamentous cells. Filamentous growth was induced for 45-150 min. Each symbol is median cell D_eff_ (*n* = 20 cells; 10-350 trajectories/cell). Mean filament length 12 μm; color gradient indicates length with **** < 0.0001. C) Cytoplasmic fluidity increases with filament length. Filamentous growth induced on agarose SC FBS, 45-125 min. Green squares are median cell D_eff_ and bars SEM. Values are grouped in 2 μm filament length bins (red to blue gradient, with color indicating length as above; *n* = 6 – 26 cells; 1700 – 2600 trajectories). Red symbol is budding cells (*n* = 67; 2500 trajectories). The solid black line is a fit using *eq* 2 using overall cell volume; r^2^ = 0.95. D) Simulation of GEM diffusion upon cell volume increase. Particle D_eff_ in a cylindrical or spherical compartment, with ribosome crowders 20% at initial cell volume 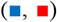. Simulations with excluded compartments (*e.g*. vacuoles;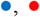), further reduced accessible volume from 80% 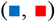 to 20% 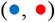. Values are D_eff_ means (6000 particle simulations) carried out 10 times. Data fit with an exponential plateau equation, r^2^ > 0.99, 95% confidence levels. E) Ribosome concentration is reduced in filamentous cells. Particle matching (cryo-EM 2DTM) of 60S ribosomal subunits in indicated cells (filament length assessed by SEM). F) Ribosome density decreases with filament length. Ribosomes concentration (*n* = 8 budding and 5 filamentous cells) determined by cryo-EM 2DTM, small symbols values per ROI, large symbols median of each cell and symbol color corresponds to filament length. Fit of medians to a straight line, r^2^ = 0.77 with 95% confidence levels; slope different from 0, *p* < 0.0001.

After 1-2 hr incubation in serum, filaments formed with an average length of 12 μm (range of 2 – 22 μm). We observed a striking increase in GEM D_eff_ of ∼2-fold in these filamentous cells, which corresponds to a substantial fluidization of the cytoplasm (Fig. 1B, S3). We speculated that some of the GEM D_eff_ variation in filamentous cells could be due to the different filament lengths. Hence, we examined whether cytoplasmic diffusivity scaled with filament length, which is directly proportional to cell volume. Figure 1C shows a correlation between filament length and cytoplasmic diffusivity; when values were grouped in 2 μm filament length bins we observe a strong positive correlation between D_eff_ and filament length, with a Pearson coefficient of 0.93. The median D_eff_ in the mother and filament compartments was examined and there was a small, but significant decrease in the filament, compared to mother compartment and whole cell (Fig. S4A). This is consistent with changes in the surface to volume, as shown in our diffusion simulations (Fig. S5, compare red-sphere to blue-cylinder bars).

Cytoplasmic fluidization scaling with increased cell volume suggests that during hyphal morphogenesis there is substantial dilution of a molecular crowder, such as ribosomes. To test this hypothesis, we simulated the diffusion of mesoscale particles in spherical and cylindrical cell geometries, with an equivalent initial crowder/cell volume of 14,000 ribosomes/μm^3^ cytoplasm, based on values from *S. cerevisiae*, which accounts for 20% of the initial cytoplasmic volume^6^. We also investigated the effect of the addition of large intracellular excluded volumes (analogous to vacuoles in volume) on the simulation of ribosome D_eff_ (Fig. S5). As expected, we found that in contrast to the small effects of cell geometry, the addition of inaccessible space dramatically reduced D_eff_ (between 50-70%, depending on geometry). Note that the increase in cell volume during morphogenesis is greater than that from solely cell geometry changes, as the vacuole, which is GEM inaccessible, is localized predominantly to the mother cell portion (Fig. S1B)^9,10^. Therefore, we simulated the effect of doubling the cell volume, which occurs approximately every hour with filament elongation, maintaining the amount of cytoplasmic crowder. Figure 1D shows that, in such a simulation (irrespective of geometry), we also observe a significant increase in D_eff_ (∼ 2-fold), similar to the experimental data (Fig. 1C). These results are consistent with a striking dilution of a cytoplasmic crowder - on the order of 4-5-fold - due to both increased overall cell volume and accessible volume in the filament, as a result of less vacuoles (Fig S6). These simulations reveal that the decreased crowding could be due to a combination of a decrease in ribosome biogenesis, and an increase in accessible volume, both of which decrease crowding.

Hence, we tested this theoretically using the simplified formalism that describes diffusion with respect to energy transfer in polydispersed mixtures (*eq 2*)^8^ to fit our data and derive the ribosome concentration, assuming they were the main crowding agent. We assumed that during filament growth, ribosomes are only diluted (*i.e*. no new synthesis). Fitting the experimental data using this equation allowed us to extract the initial concentration of ribosomes in the mother cell compartment, *i.e*. 24,000 ± 300 ribosomes/μm^3^ of cytoplasmic volume (Fig. 1C, predicted). In addition, when we took into account the fraction of GEM accessible volume (Fig. S6), in *eq 2*, we still observed a good fit to the experimental data, yielding a ribosome concentration (*c*_ribo_) of 20,000 ± 700/μm^3^ (Fig. S7), which is somewhat higher than that determined in *S. cerevisiae*^6^. The lower D_eff_ in *C. albicans* suggests that budding cells are more crowded in this fungal pathogen compared to *S. cerevisiae*.

As ribosomes are likely to be the predominant cytoplasmic crowder, we used liquid-chromatography tandem mass spectrometry (LC-MS/MS) to determine the relative abundance of ribosomal proteins, which decreased upon filamentation, compared to budding cells (Fig. S8). This is in agreement with the decreased (∼2-fold) levels of ribosomal RNA reported in *C. albicans* filamentous cells^11^, as well as a substantial decrease in the transcripts of many ribosomal proteins upon filamentation^12^. To directly assess ribosome levels during morphogenesis we carried out cryogenic electron microscopy (cryo-EM) coupled with 2D template-matching (2DTM) to identify 60S ribosomes^13,14^ (Figure S9, S10). Figure 1E shows that in filamentous cells there is a marked decrease in ribosome concentration, which is filament length dependent (Fig. 1F). Together, our results reveal that changes in crowding at the mesoscale occurs, in part, *via* ribosomes, although we cannot exclude the possibility that there is also a decrease in the concentration of larger, slowly diffusing polysomes^15^ and/or changes in cytoplasmic viscosity. The increased GEM D_eff_ in the filamentous cells could also be the result of decreased turgor pressure, hence we examined how mesoscale cytoplasmic diffusivity was affected by changes in external osmolarity using sorbitol. At all sorbitol concentrations examined we observed an increased GEM D_eff_ in filamentous compared to budding cells (Fig. 2A). Furthermore, a higher concentration of sorbitol is required to fully abolish GEM movement in filamentous cells, suggesting that molecular crowding in such cells is reduced (Fig. S11A, B).

**Figure 2.**
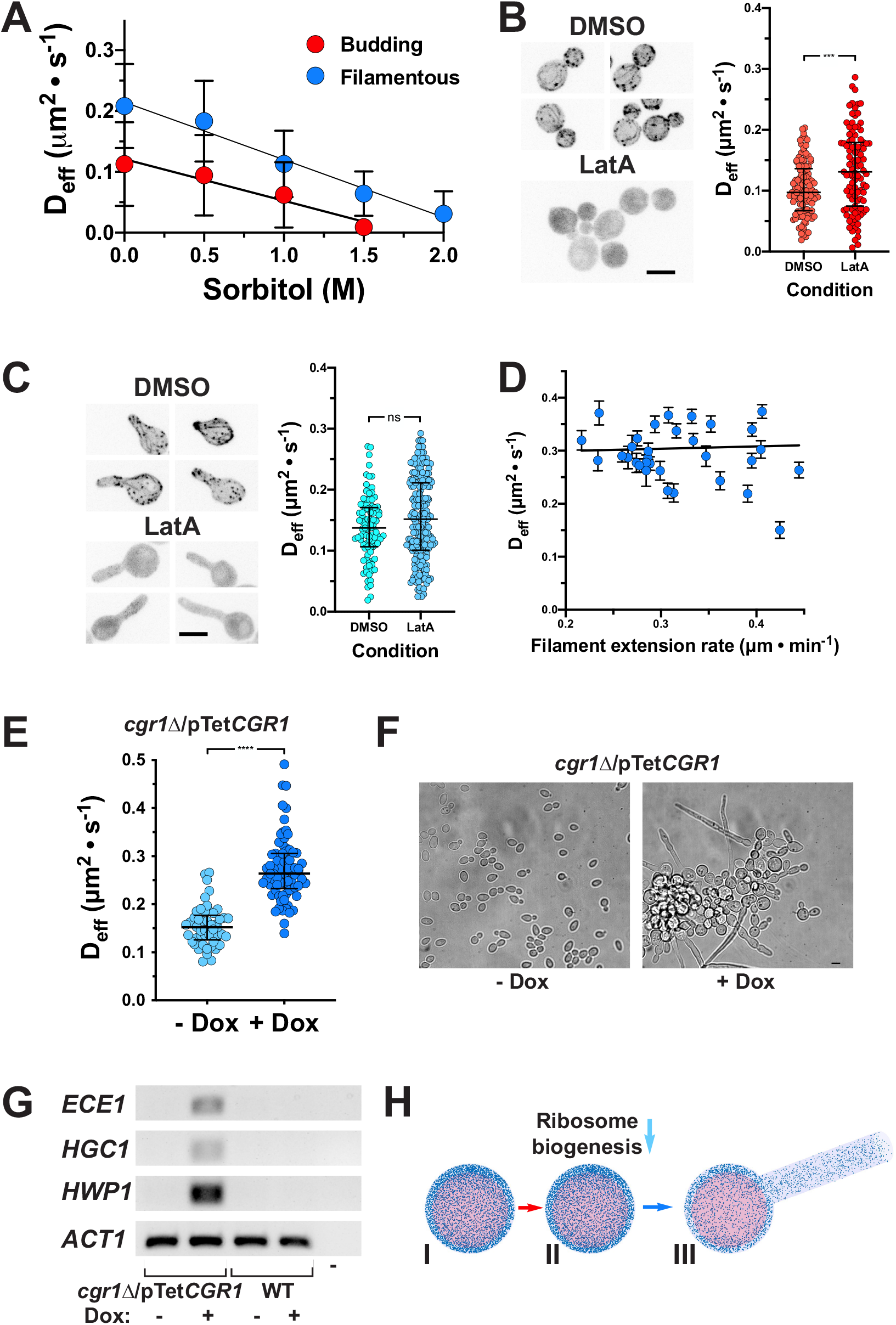
Ribosome reduction increases cytoplasmic fluidity and results in filamentous growth. A) Cytoplasmic fluidity is higher in filamentous cells, at different sorbitol concentrations. Symbols are the mean of 20-80 cells (11 - 200 trajectories/cell); bars are SD. Mean filament length was 7 ± 3 μm. B) Disruption of actin cytoskeleton in budding cells increases cytoplasmic fluidity. Left: Max projections of cells with LifeAct-RFP, with or without LatA. Right: symbol is median cell D_eff_ (*n* ∼ 100; 10-180 trajectories/cell), *** = 0.0005. C) Disruption of actin cytoskeleton in filamentous cells does not affect cytoplasmic fluidity. Left: Max projections of cells induced for 45 min with or without LatA. Right: symbol is the median cell D_eff_ (*n =* 100-200; 15-150 trajectories/cell) of filamentous cells with or without LatA, ns not significant. D) Cytoplasmic fluidity in filamentous cells is independent of growth rate. Cells preincubated on agarose SC FBS, 40 min and followed for 60 min (images every 15-20 min). Each symbol is the median cell D_eff_ with SEM and cell extension rate. Mean filament length was 17 ± 4 μm. E) *CGR1* repression increases cytoplasmic fluidity. Each symbol is the median cell D_eff_ (*n* = 50-75; 30-250 trajectories/cell); cells grown with or without Dox, **** < 0.0001. F) *CGR1* repression promotes filamentous growth. Representative cells with or without Dox. G) Hyphal-specific genes are induced upon *CGR1* repression. *ECE1, HGC1* and *HWP1* transcripts in indicated strains, with or without Dox; *ACT1* as internal control. H) Schematic of ribosome biogenesis inhibition triggering to filamentation. Red arrow indicates serum addition, which inhibits ribosome biogenesis (light blue arrow), leading to ribosome dilution upon growth (dark blue arrow). Blue dots are ribosomes and pink sphere vacuole compartment, inaccessible to ribosomes.

We also examined whether mesoscale cytoplasmic diffusivity was affected by depolymerization of the actin cytoskeleton, as it was shown to decrease cytoplasmic diffusivity in *S. cerevisiae*^6^, yet had no effect in *S. pombe*^16^. Figures 2B and S12 show that in budding *C. albicans* cells the GEM D_eff_ increased

∼35% upon Latrunculin A (LatA) disruption of F-actin. In contrast, the same treatment had a substantially smaller effect on the GEM cytoplasmic dynamics in filamentous cells (Fig. 2C and S12). However, in LatA treated filamentous cells there was no longer a difference between the GEM D_eff_ in the filament compartment and that of the whole cell (Fig. S4B), indicating that the actin cytoskeleton, rather than solely cell geometry, is important for the small difference in the cylindrical filament (Fig. S5). Overall, these results indicate that actin is more critical for cytoplasmic crowding in budding cells compared to hyphal cells, in part due to the decreased accessible cytoplasm volume in budding cells (Fig. S6). As LatA also blocks growth, these results also reveal that growth *per se* does not substantially contribute to cytoplasmic mesoscale crowding in filamentous cells. This is further confirmed in Figure 2D, which shows that over a 2-fold range of filament extension rates, GEM D_eff_ was essentially constant.

The increase in cytoplasmic fluidity, as a result of the decreased ribosome concentration upon filament elongation, suggested that inhibition of ribosome biogenesis may be important for filamentous growth. This would lead to dilution of ribosome crowders upon subsequent growth. To investigate the role of ribosome biogenesis in morphogenesis, we generated a mutant defective in the former process. *CGR1* encodes a protein that is critical for the processing of pre-rRNA in *S. cerevisiae*^17^, in particular rRNA for the 60S ribosome subunit. Addition of doxycycline (Dox) to a *C. albicans* strain in which the sole copy of *CGR1* is under the control of the Tet-repressible promoter, resulted in complete repression of *CGR1* mRNA transcript (Fig. S13A) and in slow growth (Fig. S13B). Analyses of rRNA levels revealed a decrease in 28S and 18S rRNA upon *CGR1* repression (Fig. S13C) and a substantial (∼1.7-fold) increase in GEM D_eff_ (Fig 2E). Strikingly, repression of *CGR1* resulted in some filamentous cells in the absence of serum, which was not observed in the absence of Dox nor in wild-type cells (Fig. 2F). Furthermore, Figure 2G shows a dramatic induction of the hyphal specific genes *e.g*. those encoding the G1 cyclin, *HGC1*, the candidalysin toxin, *ECE1*, and the hyphal cell wall glycoprotein, *HWP1*, in the *cgr1* mutant in the presence of Dox. Similarly, inhibition of the Tor kinase by rapamycin, which has been shown to reduce the number of ribosomes in *S. cerevisiae*, resulted in hyphal specific gene induction in *C. albicans*^18^. Together these results reveal that inhibition of ribosomal biogenesis is important for the yeast to hyphal morphogenetic transition and that, upon filament elongation, dilution of ribosomes leads to a substantial reduction in cytoplasmic crowding.

In *S. cerevisiae*, a sizeable portion of ribosomes (at least 25%) do not contribute to translation^19^. Intriguingly, while *C. albicans* filamentous growth (0.3 μm/min extension) is similar with respect to volume increase, when compared to budding growth (doubling every 90 min), genome dilution upon DNA replication arrest in *E. coli* was reported to result in a decrease in active ribosomes and sub-optimal growth^20^. Increased cytoplasmic fluidity has been shown to increase cytoskeleton polymerization and depolymerization rates in fission yeast^5^, suggesting that decreased molecular crowding at the mesoscale could also be beneficial for *C. albicans* filamentous growth. In summary, our results reveal a striking change in cytoplasmic molecular crowding during morphogenesis (Fig. 2H), which is critical for the virulence of this human fungal pathogen, as a result of dilution of ribosomes subsequent to inhibition of their biogenesis. Our results suggest that changes in cytoplasmic diffusion at the mesoscale, by tuning ribosome numbers, are intimately associated with filamentous growth.

## MATERIAL AND METHODS

### Strains and media

Strains used in this study are listed in Table S1. For transformation, strains were grown in YEPD (yeast extract, peptone, dextrose) supplemented with Uridine (80 μg/ml) at 30ºC. Cells were grown in YEPD medium, supplemented with Uridine at 30ºC for budding growth. For filament induction cells, cells were either grown in YEPD liquid media with 50% fetal bovine serum (FBS; PAN Biotech) or on agarose pads with 75% FBS in synthetic complete (SC) media, both at 37ºC. In all experiments that involved comparison with filamentous cells, budding cells were briefly incubated in the same media at 37ºC prior to imaging at the same temperature. For doxycycline (Dox) gene repression, YEPD was supplemented with 5 μg/ml Dox. For sorbitol experiments, budding and filamentous cells were incubated with the indicated final concentrations of sorbitol for 5 min prior to imaging at 37ºC. For Latrunculin A (LatA) actin depolymerizaton, cells were incubated with either 200 or 400 μM LatA, for budding and filamentous cells, respectively, during 15 min prior to imaging.

The oligonucleotides used in this study are listed in Table S2. The PfV gene, which encodes the subunits that comprise a 40-nm genetically encoded multimeric (GEM) nanoparticle^1^ was codon optimized for *C. albicans* and synthesized (BaseClear, Netherlands). The CaPfV gene was cloned from the pUC57 vector into the pFA-GFPγ-URA3 plasmid using PstI sites, resulting in the pFA-CaPfV-GFPγ-URA3 plasmid. The *URA3* marker was replaced with either *CdHIS1* and *ARG4* markers, using unique AscI and PmeI restriction sites, resulting in pFA-CaPfV-GFPγ-CdHIS1 and pFA-CaPfV-GFPγ-ARG4. The GFPγ was then mutated using site directed mutagenesis to a monomeric version using primers CaGFPγpA206K-BamHI/CaGFPγmA206K-BamHI, yielding pFA-CaPfV-GFPγ^A206K^-CdHIS1 and pFA-CaPfV-GFPγ^A206K^-ARG4. Subsequently these two plasmids were used as a template to PCR amplify the CaPfV-GFPγ^A206K^-CdHIS1 and pFA-CaPfV-GFPγ^A206K^-ARG4 cassettes for integration behind the endogenous *ADH1* promoter by using primers ADH1p-PfVp and CaADH1KixFP_S2. The tetracycline repressible *cgr1*Δ/pTet*CGR1* strain was constructed from PY173, a derivative of BWP17 containing the tetracycline-regulatable transactivator TetR-ScHAP4AD, as described^2^. To visualize F-actin, a LifeAct reporter was used which was derived from the plasmid pYGS974 (TEF1ΔLifeAct-GFP-HIS1/pJET)^3^, in which GFP was replaced by mScarlet using primers NotITef1pLifeActS1 and CamScarletmAscI, resulting in pTEF1p-LifeAct-mScarlet-HIS1. This plasmid was digested with NotI/XbaI and integrated by homologous recombination in the *TEF1* locus.

### Microscopy, sample preparation and image analysis

#### Light microscopy

The GEM nanoparticles were imaged, using TIRF on a Nikon Ti eclipse inverted microscope (Nikon France S.A.S., Champigny-sur-Marne, France) equipped with an iLas^2^ scan head (Roper scientific, Evry, France), an iXon 888 EMCCD camera (Andor technology, Belfast, UK) and a 100x CFI-APO-TIRF oil NA 1.49 objective. The laser illumination was with a 488 nm diode laser, with an intensity ranging between 10-60% and a TIRF angle of 56.24º. Images were acquired as a stream of 300-600 images in a single Z plane, with a 30 msec image acquisiton and readout time, unless otherwise indicated. Temperature was controlled with an Okolab incubator (Ottaviano, Italy) at 37ºC, unless otherwise indicated. The LifeAct reporter was imaged using the spinning-disk confocal modality on the above described microscope, equipped with a Yokogawa CSU-W1 (Yokogawa Electric Corporation, Tokyo, Japan), and using a 561 nm diode laser. Multi-positions were acquired with a motorized XYZ stage. The effective diffusion of the GEM particles was calculated as described^1^, using the plugin Mosaic^4^ in Fiji (version 1.54f) in a Windows 10/Intel computer and Matlab (version R2023a). Matlab was also used to generate the plots of all trajectories using a colormap, and to split the filamentous cells in order to analyze mother cell vs filament effective diffusion (scripts available upon request). Extension rates were calculated as a function of filament length (measured with Fiji) per time. Scale bar is 5 μm.

#### Electron microscopy

Cryoplunging was carried out essentially as described^5^. Au grids (200 mesh) with a 2/2 silicone oxide support film were glow-discharged on both sides for 45 sec at 20 mA. Glycerol (5% final concentration) was added to cells and then immediately 3 μl (0.2 OD_600_) cell suspension was applied to the support film side of grids, blotted for 10 sec and then frozen in liquid ethane using a cryoplunger (GP2 Leica, Wetzlar, Germany) and frozen. FIB-milling was carried out using an Aquilos 2 FIB/SEM (Thermo Fisher, Waltham, MA) with a stage cooled to < -190°C in a 35° AutoGrid sample holder. Grids were sputter-coated with metallic Pt and then coated with organo-Pt essentially as described^5^. An overview of the grid was created by montaging SEM images and isolated cells or cell clusters at the center of grid squares were selected for FIB-milling. Lamellae were generated automatically using the AutoTEM software (Thermo Fisher, Waltham, MA), with the following protocol: rough milling 1: 1 nA; medium milling 2: 0.3 nA (1.0° overtilt); fine milling 0.1 nA (0.5° overtilt); finer milling 0.1 nA (0.5° overtilt); lamella polishing: 50 and 30 pA, 0.2°overtilt resulting in 150 nm thick lamellae which were subsequently sputter-coated with Pt for 5 sec at a current of 5mA.

FIB-milled grids were imaged in a Titan Krios TEM (Thermo Fisher, Waltham, MA) operated at 300 keV and equipped with a BioQuantum energy filter (Gatan, Pleasanton, CA) and K3 camera (Gatan, Pleasanton, CA). The instrument was controlled using SerialEM^6^. Individual lamellae were manually centered in the microscope and then moved to a position 60 μm below the eucentric height to achieve fringe-free illumination. The stage was tiled 15º to compensate for the milling angle and overview images were obtained with a pixel size of 76.8Å. Individual cells were annotated in these overview images using napari^7^ and high-resolution montages were obtained for these cells using DeCo-LACE acquisition scripts^8^. The physical pixel size in high-resolution exposures was 1.05Å, defocus was maintained at 1 μm, and the total exposure was 30 e/Å^2^. The exposures were dose-fractionated into 30 frames.

Movies were imported into the cisTEM software package^9^ and motion-corrected using a custom version of unblur^10^ as described^8^, and binned to a final pixel size of 2.0 Å. CTF parameters and sample thickness were estimated using CTFFIND5^11^. The structure of the vacant *C. albicans* 80S ribosome^12^ (PDB code: 7PZY) was modified by deleting subunits corresponding to the 40S subunit and the resulting 60S structure was converted to a density map at 2.0 Å pixel size using the simulate program in cisTEM^13^ using a B-factor scaling of 2. 2D template matching of individual exposures was performed using the GPU-accelerated version of the match_template program in the cisTEM suite^14^. Rotation angles were search using a 2 in-plane and 3 out-of-plane step-size and defocus values were searched with a 240 nm slab at a 40 nm step size. Image data and template matching results were montaged together as described^8^.

The number of 60S ribosome subunits per imaged area was determined by manually segmenting cytoplasmic areas from the montaged cryo-EM images (Fig. 1F, S9 and S10) and dividing the number of detections within the segmentation by the segmented area. The imaged volume was calculated by fitting the thickness of individual exposures, as estimated by CTFFIND5, to a 2-dimensional cubic B-spline model with 3 knots^19^ and integrating the estimated thickness at every pixel of the segmented area. The variation of this measurement was estimated by repeating this calculation in 50 random square areas with a side-length of 200 nm within the segmented area.

### Modeling cytoplasmic diffusion

#### Predictive equations

A simplified formalism^15^ was used to describe diffusion with respect to energy transfer in polydispersed mixtures to determine the fit to our experimental data and from these equations (*eq 1* and *2*) we derived the number of ribosome crowders:

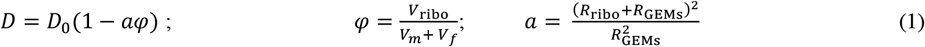

With *V*_*i*_ being the volume of the ribosomes (ribo), the mother part of the cell (m) and the filament (f). *D*_*0*_ is the diffusion coefficient of GEMs in the absence of ribosomes, but with other crowders in the cell. To avoid this unknown, we define 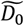 as the diffusion coefficient of GEMs when the there is no filament (about 0.1 μm^2^/s in our case). This results in:

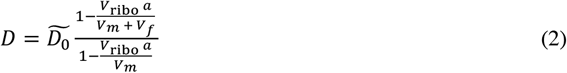

We used *eq* (2) to fit our data. Since we assume that no more ribosomes are produced during the growth of the filament, the total volume of ribosomes is fixed by the initial condition of the volume of the mother compartment, and is equal to *V*_*ribo*_ = *c*_*ribo*_ × *v*_*ribo*_ × *V*_*m*_ where *v*_*ribo*_ = 4/3π*R*_*ribo*_^3^ is the volume of 1 ribosome, assuming a spherical geometry. The only unknown in *eq* (2) is thus *c*_*ribo*_.

#### Simulations

A Matlab-based model (version R2023a) was developed to simulate particle diffusion within spherical and cylindrical boundaries, which we refer to as DiffSim. Particles of radii and concentration similar to that of ribosomes (r = 14 nm and 11,100/μm^3^) were randomly localized within these geometries, together with larger spherical crowders (similar in size to a vacuole taking up 60% of cell volume) which were immobile. To facilitate computation in this 3D simulation, cell geometries and excluded volumes were chosen accordingly. The model incorporated three primary forces affecting particle diffusion: Brownian motion and collisions with both other particles and crowders. Brownian motion was modeled using a random walk algorithm, where particles were displaced randomly at each time step. Collisions were simulated using a hard-sphere model, deflecting particles upon contact. Similarly, when particles encountered external boundaries, they were deflected back into the system. Particle positions were recorded in a matrix for subsequent analysis. The mean-squared displacement (MSD) was calculated as a function of time (with Δt being identical to GEM acquisitions) to assess the effective diffusion coefficient and its dependence on relative cell geometry and volume was determined. The effective diffusion coefficient (D_eff_) was then determined from the slope of the MSD curve using the Einstein-Smoluchowski equation. The accessible volume was determined using a custom Matlab program, which we refer to as AccessCyto, which analyzed the positions of GEM particle tracks to calculate the region of the cell that was accessible to particle movement. The accessible volume was then compared to the total cell volume, by segmenting the maximum projections, to calculate the percentage of the cell accessible to the GEM particles. Scripts available upon request.

#### Proteomics

Sample preparation was essentially as described^16^. Quality control samples to monitor LC-MS performance were created from pooling small aliquots of all samples. Peptide quantities were estimated *via* Quantitative Fluorometric Peptide Assay (Pierce). LC-MS based proteomic data acquisition was performed as described^17^. In brief, samples were injected on a ACQUITY M-Class HPLC (Waters) connected to a ZenoTOF 7600 mass spectrometer with an Optiflow source (SCIEX), separated on a HSS T3 column (300 μm×150 mm, 1.8 μm; Waters) using a 20 min active gradient. We used a Zeno SWATH acquisition scheme with 85 variable-sized windows and 11 ms accumulation time. LC-MS raw data was processed using DIA-NN 1.8^18^. First, a spectral library was predicted including the UniProt Proteome of *C. albicans* SC5314 (https://www.uniprot.org/proteomes/UP000000559), as well as the sequence of the genetically encoded multimeric nanoparticle (see description above). For the main search, we enabled tryptic digestion allowing for one missed cleavage, no variable modification, N-terminal Methionine excision and carbamidomethylation as fixed modification of Cysteines. Mass accuracies were set to fall within 20 ppm and match-between-runs was enabled with protein inference on Protein level. The obtained report was processed using Python 3.9 with the pandas (1.4.3), NumPy (1.23.0) and Seaborn (0.11.2) packages. Data was filtered to less or equal than 1% FDR concerning Global.Q.Value, as well as PG.Q.Value and Lib.PG.Q.Value. Prior to plotting, protein group intensities were sample-wise median normalized in two steps: first, subtracting the sample median in log2-space, then subtracting the log2-sample median of entities not belonging to the ribosome, according to UniProt annotated protein names (*i.e*. containing “60S” or “40S, that is the 76 core ribosome proteins).

#### RNA extraction and RT-PCR

Cells were grown in YEPD media in the presence or absence of 5 μg/ml of Dox. RNA extraction and RT-PCR were carried out as described^20^. Oligonucleotide pairs ACT1.P1/ACT1.P2, CGR1.P1/CGR1.P2, ECE1.P1/ECE1.P2, HGC1.P1/HGC1.P2 and HWP1.P1/HWP1.P2 were used to amplify *ACT1, CGR1, ECE1, HGC1* and *HWP1*, respectively.

### Statistical analysis

Data were compared by the Mann-Whitney U test and where relevant the paired or unpaired t-test using GraphPad Prism (v. 8) software, with all *p* values indicated in figure legends. Unless stated otherwise medians and interquartile ranges are indicated. Pearson correlation coefficient and simple linear regression were determined using GraphPad Prism (v. 8) software.

## Acknowledgements

We thank P. Silva, H. Labbaoui and S. Bogliolo for assistance, S. Noselli and A. Hubstenberger for comments on the manuscript, the PRISM Imaging facility (B. Monterroso and S. Ben-Aicha) and the Microscopy Imaging Cytometry d’Azur (MICA) for microscopy support, the BIOINFO Bioinformatic facility (A. Fortuné) for computational support, and M. Rigney for support with cryo-EM sample preparation. Cryo-EM data were acquired at the UMass Chan Medical School Cryo-EM Core Facility. This work was supported by the CNRS, INSERM, Université Côte d’Azur, ANR (ANR-19-CE13-0004-01), EC (MSCA-ITN-2015-675407; MSCA-IF-2020-101029870; ERC-SyG-2020 951475) and FRM (SPF202309017657) grants.

## Supplementary Figure legends

**Figure S1**. Projections of GEM trajectories. Representative budding and filamentous cells with maximum projection of GEM images (A) and particle trajectories of GEMs (B) shown. False colored LUT is indicated.

**Figure S2**. GEM effective diffusion is independent of its expression level. Each symbol represents the median D_eff_ of indicated cells (*n* = 25-35 cells each condition; 25-120 trajectories per cell), expressing GEMs under the control of either the *ADH1* or the *TEF1* promoter and grown at 30ºC, with medians and interquartile range indicated; ns, not significant.

**Figure S3**. Effective diffusion of all trajectories from budding and filamentous cells. Cells (20 each) from Fig. 1B, with 900 – 3300 trajectories for each condition. Medians and interquartile range are indicated; **** < 0.0001.

**Figure S4**. GEM effective diffusion is somewhat reduced in filament compartment. A) GEM effective diffusion in filament and mother cell compartments. Left panel: median D_eff_ of cells (*n* = 74) from Fig. 2C, in the absence of latrunculin A (LatA), with trajectories in whole cell, mother compartment and filament compartment analyzed. Medians and interquartile range are indicated, with ** < 0.005 paired t-test. Right panel: effective diffusion of all trajectories from filamentous cells from Fig. 2C in the absence of LatA, with trajectories in whole cell (*n* = 11100), mother compartment (*n* = 6300) and filament compartment (*n* = 4700) analyzed. Medians and interquartile range are indicated, with *** < 0.001 and * < 0.01. B) GEM effective diffusion is similar in the mother and filament cell compartments upon disruption of the actin cytoskeleton. Left panel: median D_eff_ of cells (*n* = 74) from Fig. 2C in the presence of LatA, with all trajectories in whole cell, mother compartment and filament compartment analyzed. Medians and interquartile range are indicated with ns, not significant paired t-test. Right panel: effective diffusion of all trajectories from filamentous cells from Fig. 2C in the presence of LatA with trajectories in whole cell (*n* = 7300), mother compartment (*n* = 5000) and filament compartment analyzed (*n* = 2100). Medians and interquartile range are indicated with ns, not significant.

**Figure S5**. Simulation of GEM diffusion as a function of crowding. A) Representation of particle diffusion simulation with ribosomes (blue dots) and vacuoles (large red spheres), within the cell boundaries. B) Effect of cell geometry and excluded internal volume on particle diffusion. Ribosome crowding was initially set to 20% of the cell compartment volume, either in the presence or absence of vacuole compartment, which excluded an additional 60% of the cell. Note the surface to volume ratio increases ∼3-fold from a sphere to a cylinder of fungal cell size.

**Figure S6**. The GEM accessible volume increases significantly as the filament extends. The positions of the GEMs and the outer edge of cells used in Fig. 1C was used to calculate the relative volume of the whole cell 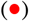 and the GEM accessible cytoplasm 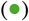. The fraction of GEM accessible volume 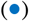 is the volume of the GEM accessible cytoplasm divided by the volume of the whole cell. Error bars indicate standard deviation, data was fit either to a straight line (r^2^ = 0.72-0.76) or one phase association (r^2^ = 0.48) for each condition with 95% confidence levels shown.

**Figure S7**. Theoretical equation for diffusion yields a good fit to the experimental data. Symbols (circles, red to blue color indicating filament length) are data (filament length bins) from Fig. 1C and grey squares/predicted line are from theoretical equation (*eq 2*), using GEM accessible volumes from the results in Fig. S6. Correlation coefficient, r^2^ = 0.83.

**Figure S8**. Relative abundance of ribosomal proteins decreases upon filamentation. Ribosomal proteins levels were determined by Liquid-chromatography tandem mass spectrometry (LC-MS/MS) from independent (n = 4) replicates, and normalized to the sample median protein levels. Control cells were grown in YEPD at 30ºC. In serum conditions, 50% FBS was added and samples were incubated at 37ºC for 0 and 90 min. Bars indicate means and ** < 0.01.

**Figure S9**. 2D Template matching of 60S ribosomal subunits in cryo-EM images of budding cells. A) Montage of cryo-EM exposures of three representative cells. B) Manual segmentation of cytoplasmic regions. C) 2DTM detections of the 60S ribosomal subunit. D) Overlays of A-C. E) Histogram of 2DTM SNR of 60S ribosomal subunit detections.

**Figure S10**. 2D Template matching of 60S ribosomal subunits in cryo-EM images of filamentous cells. A) Images of three representative cells taken by the focused ion beam prior to milling. Length of filaments was measured using the image viewer of the AutoTEM software. B) Montage of cryo-EM exposures of representative cells. C) Manual segmentation of cytoplasmic regions. D) 2DTM detections of the 60S ribosomal subunit. E) Overlays of B-D. F) Histogram of 2DTM SNR of 60S ribosomal subunit detections.

**Figure S11**. GEM effective diffusion is reduced with increasing sorbitol concentrations. A) Increasing sorbitol concentration decreases GEM effective diffusion in budding cells. Median D_eff_ of cells (*n* = 20-80) from Fig. 2A, with 11 - 200 trajectories per cell (left), and effective diffusion of all trajectories from budding cells in Fig. 2A, with 600-4000 trajectories (right), as a function of sorbitol concentration. Medians and interquartile range are indicated. B) A higher concentration of sorbitol is required to fully abolish GEM dynamics in filamentous cells, compared to budding cells. Median D_eff_ of cells (*n* = 20-80) from Fig. 2A with 11 - 200 trajectories per cell (left), and effective diffusion of all trajectories from filamentous cells from Fig. 2A, with 600-4000 trajectories (right), as a function of sorbitol concentration. Medians and interquartile range are indicated.

**Figure S12**. The actin cytoskeleton restricts GEM diffusion in budding cells. Left panel: effective diffusion of all trajectories from budding cells in the presence and absence (DMSO) of LatA. Cells from Fig. 2B, with 5800-7200 trajectories for each condition. Medians and interquartile range are indicated; **** < 0.0001. Right panel: effective diffusion of all trajectories from filamentous cells in the presence and absence of LatA. Cells from Fig. 2C, with 13600-21000 trajectories for each condition. Medians and interquartile range are indicated; **** < 0.0001.

**Figure S13**. Repression of *CGR1* results in slow growth and reduction of rRNA levels. A) *CGR1* is fully repressed in doxycycline. The transcript level of *CGR1* was determined in *cgr1*Δ*/*pTet*CGR1* and wild-type strains, grown in the presence or absence of 5 μg/ml Dox by RT-PCR with *ACT1* used as an internal control. B) Repression of *CGR1* results in slow growth. Indicated strains were incubated with or without 5 μg/ml Dox on rich media containing agar for 2 days. C) Repression of *CGR1* results in a decrease in rRNA. Total RNA was isolated from the indicated strains.

**Table S1.**
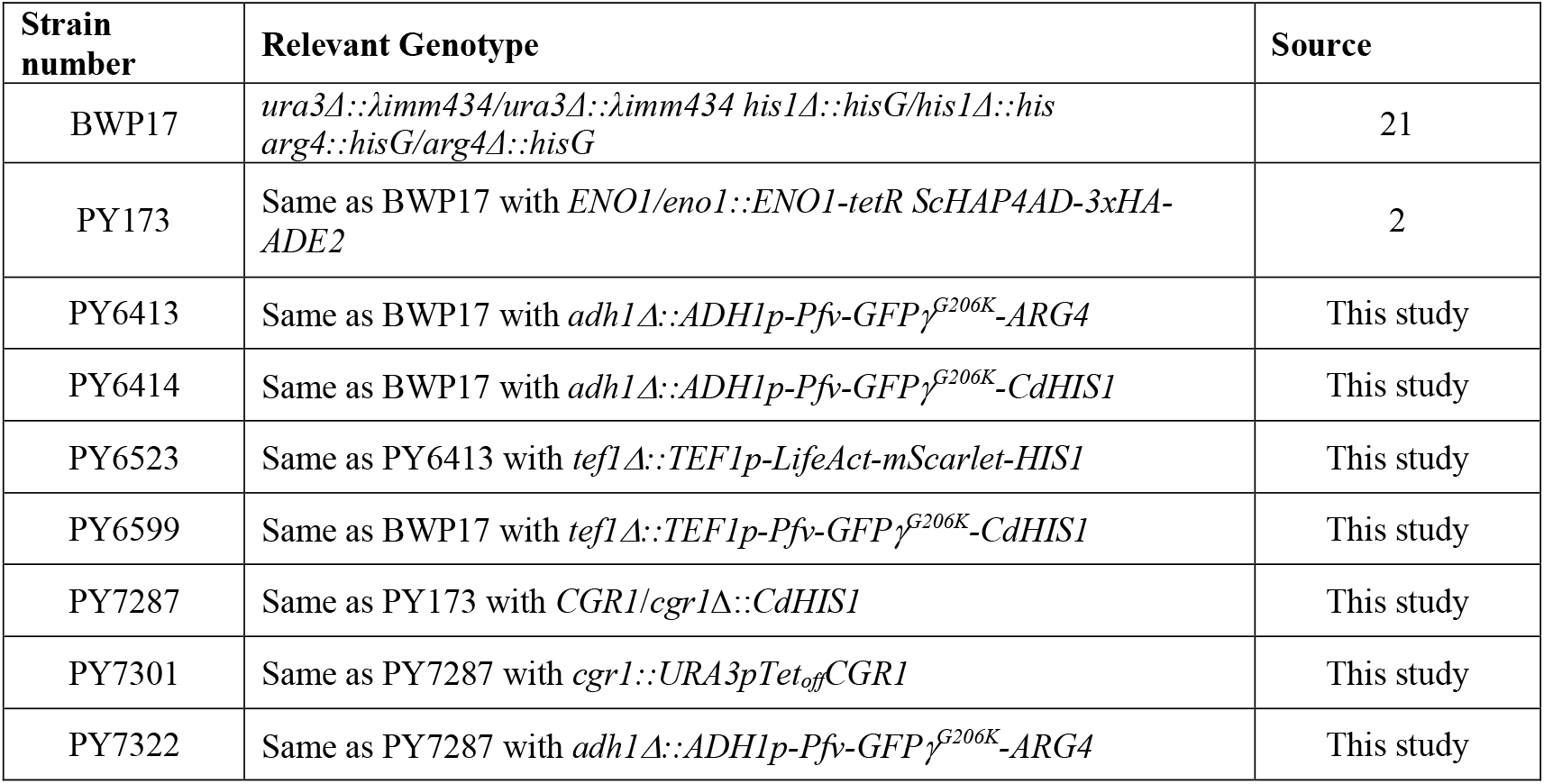
Strains used in this study.

**Table S2.**
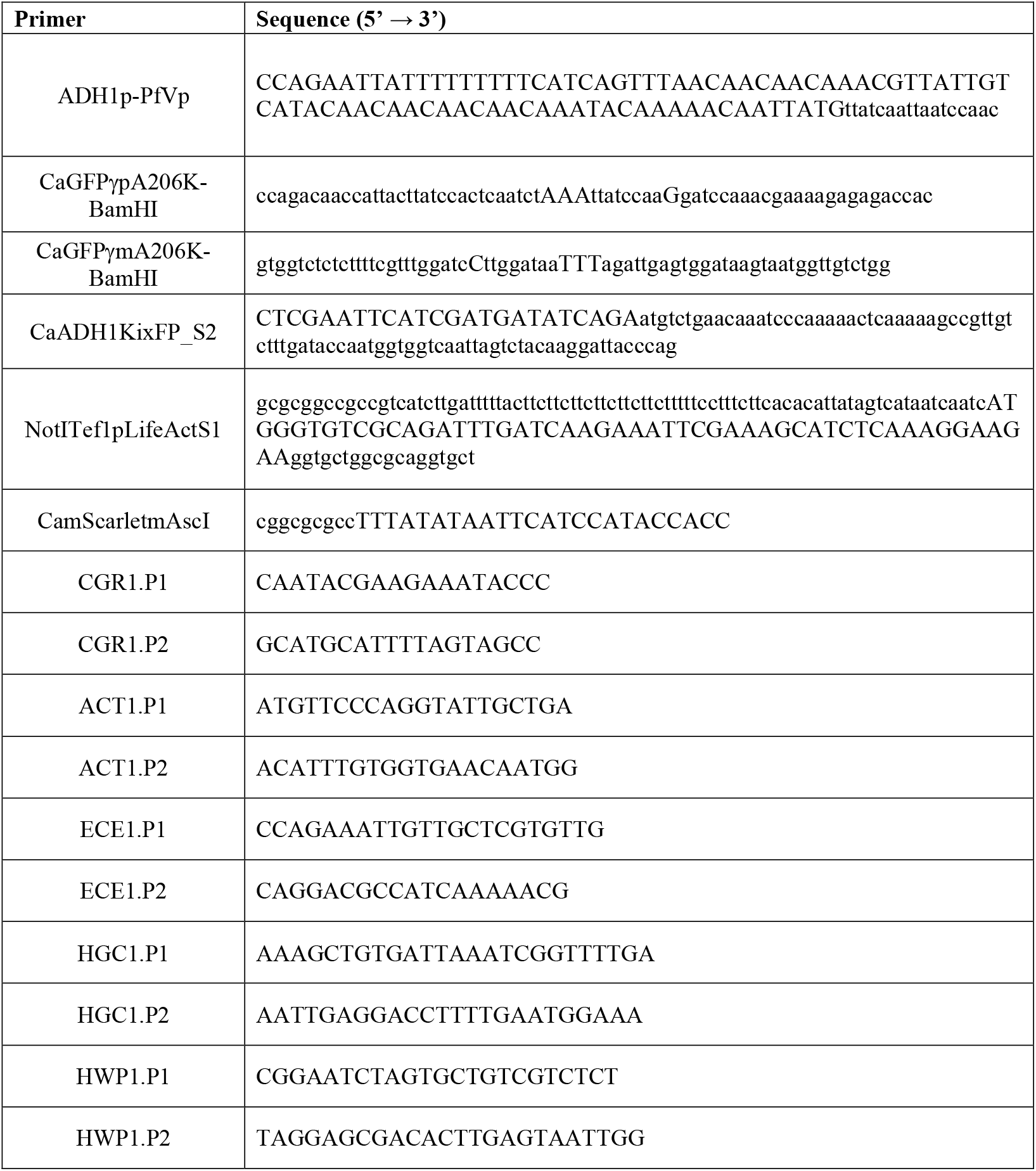
Oligonucleotides used in this study.

