## Supplemental Figures 1-13 for "Dynamic cytoplasmic fluidity during morphogenesis in a human fungal pathogen"

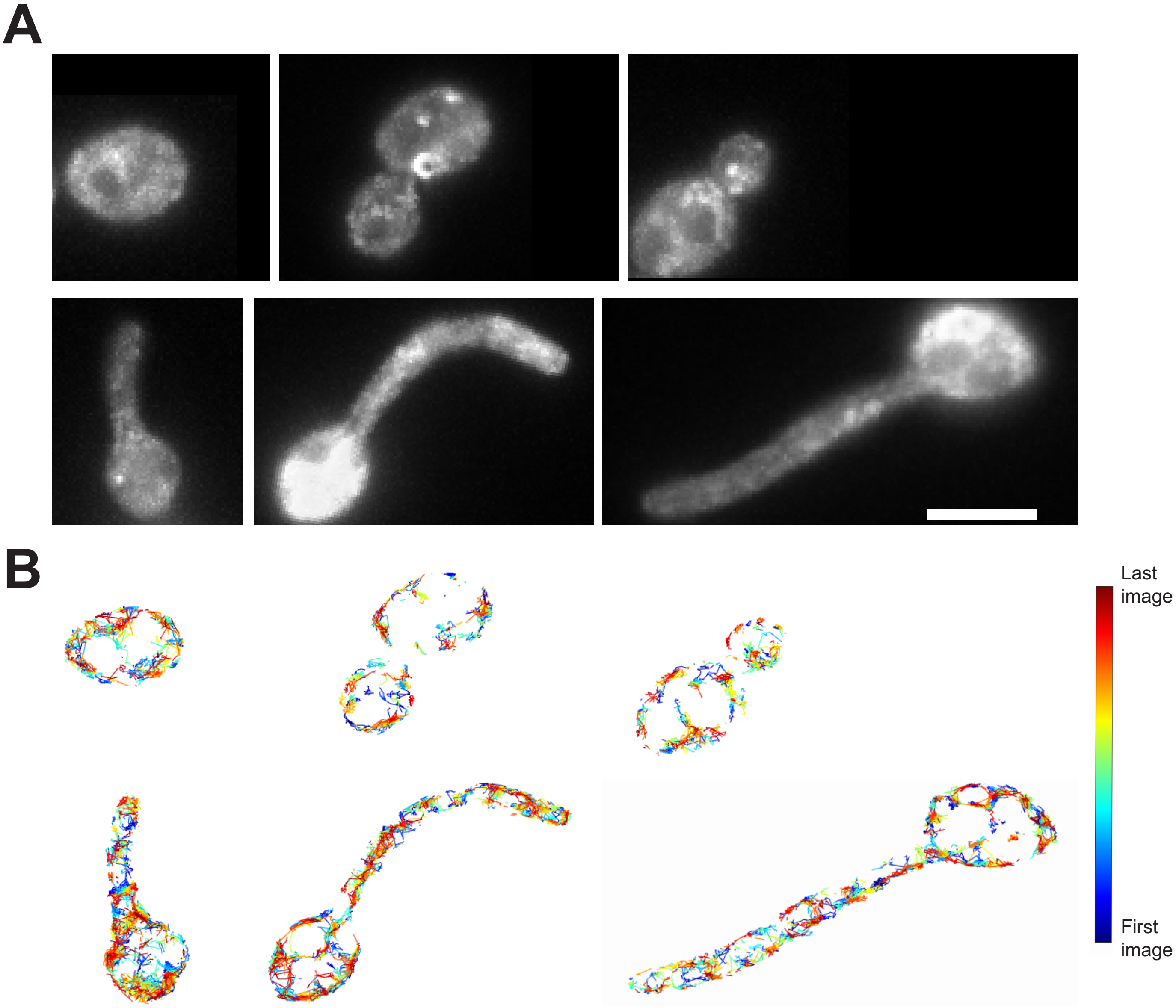


**Figure S1**

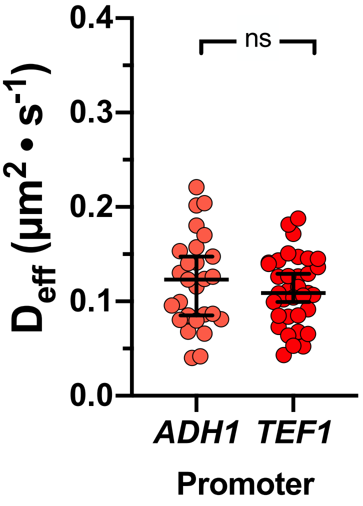


**Figure S2**

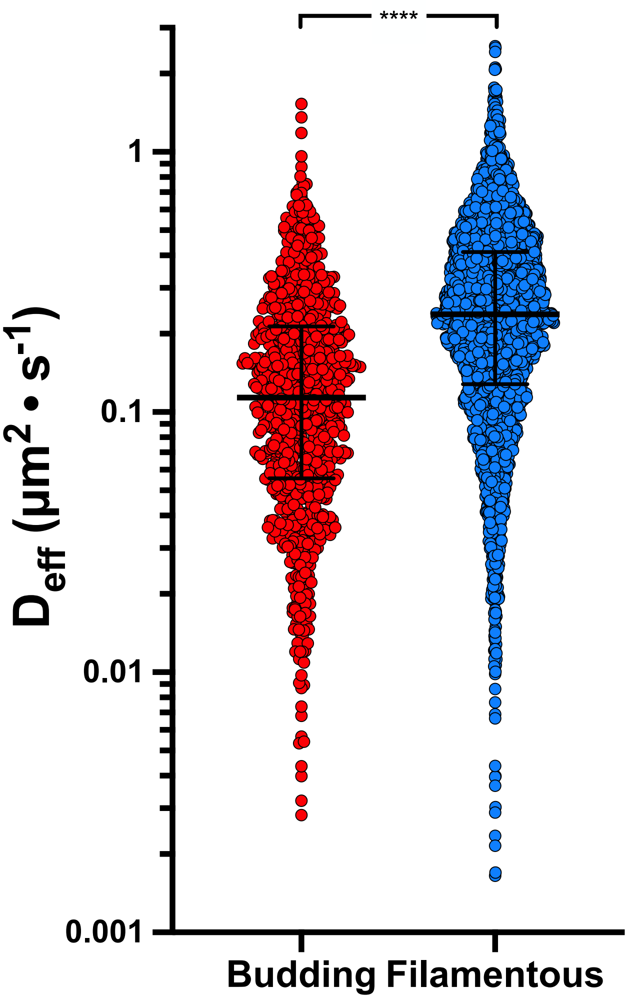


Figure S3

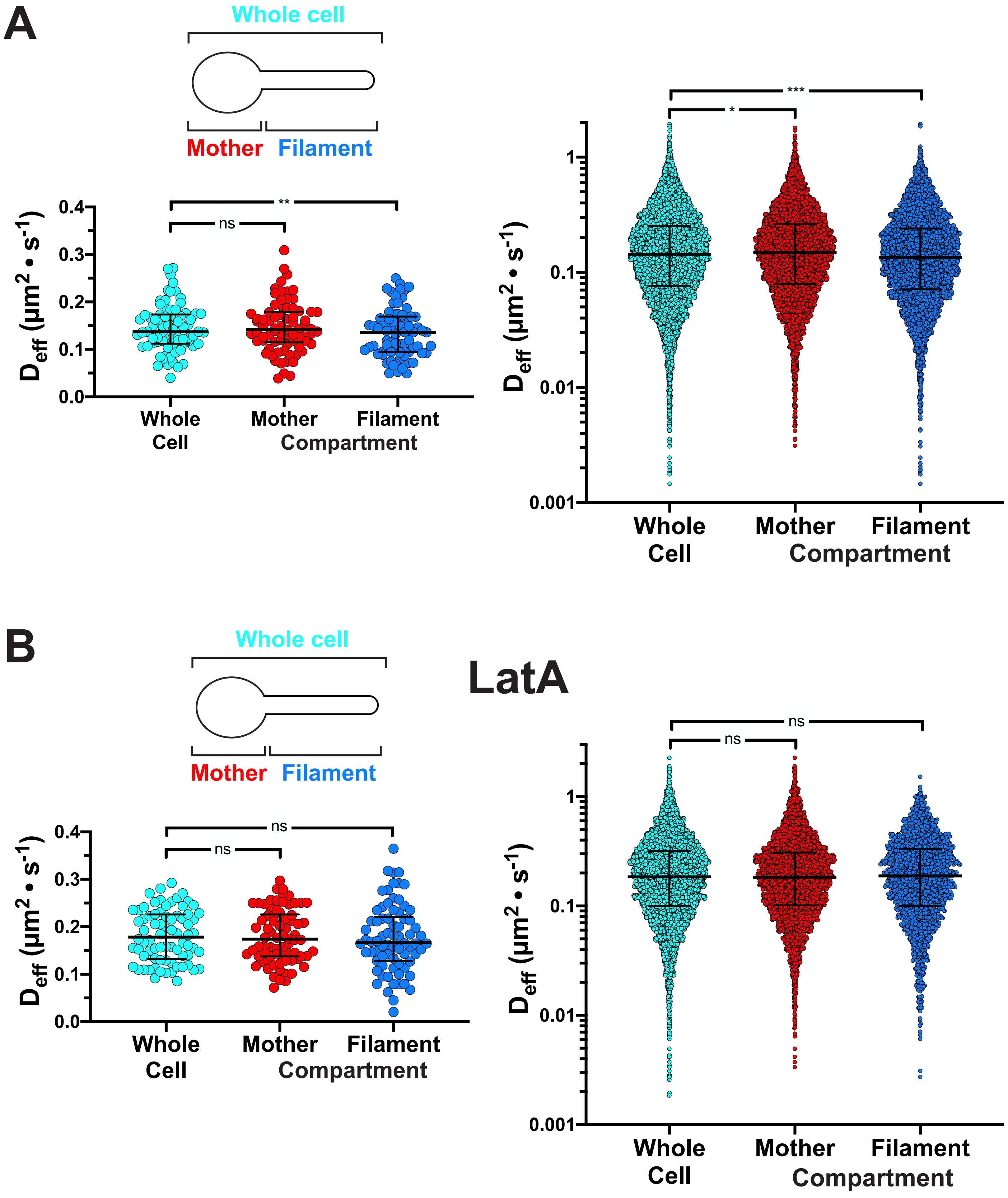
A
Whole cell
Mother
Filament
D$_{eff}$ (µm$^2$ • s$^{-1}$)
0.4
0.3
0.2
0.1
0.0
**
ns
***
*
1
0.1
0.01
0.001
Whole Cell
Mother
Filament
Compartment
B
Whole cell
Mother
Filament
LatA
ns
ns
ns
ns
Whole Cell
Mother
Filament
Compartment

Figure S4

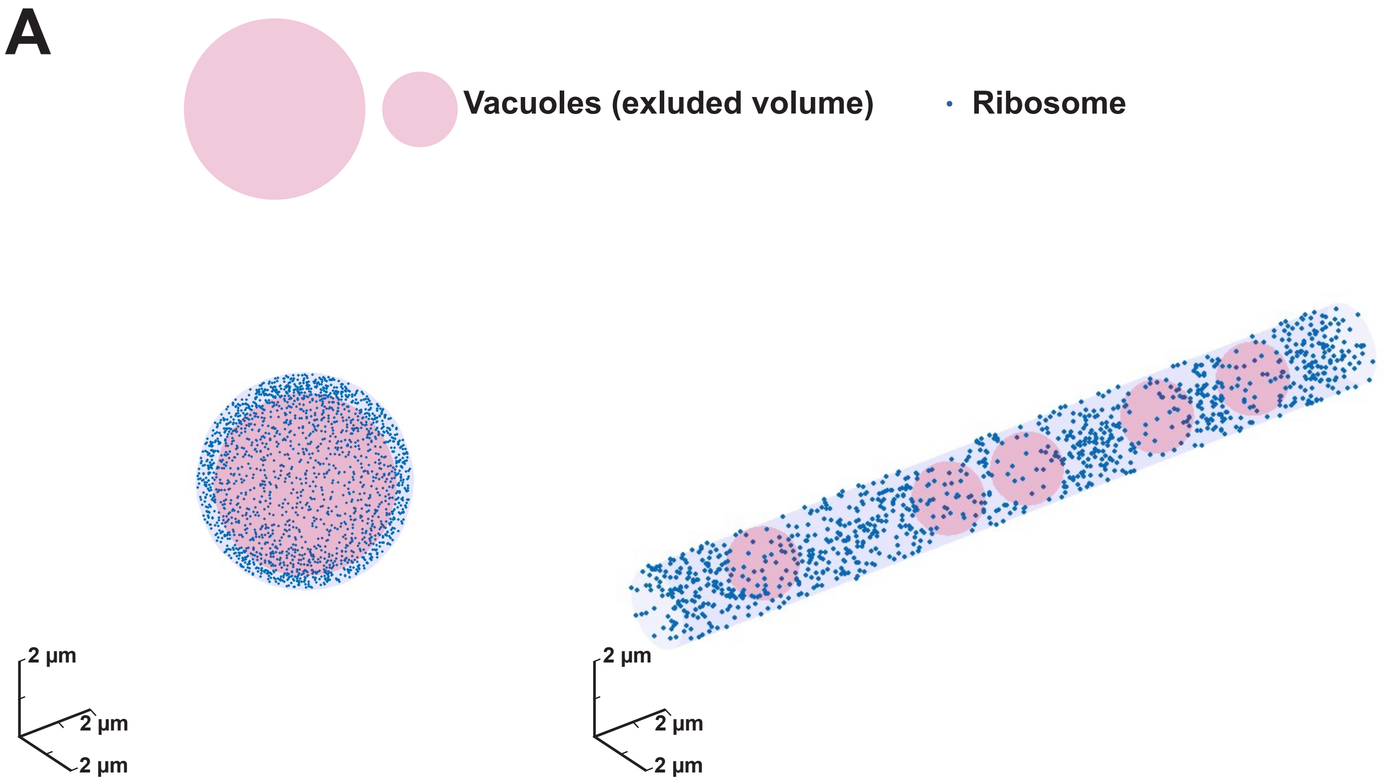
A
Vacuoles (exluded volume)
Ribosome
2 µm
2 µm
2 µm
2 µm
2 µm
2 µm

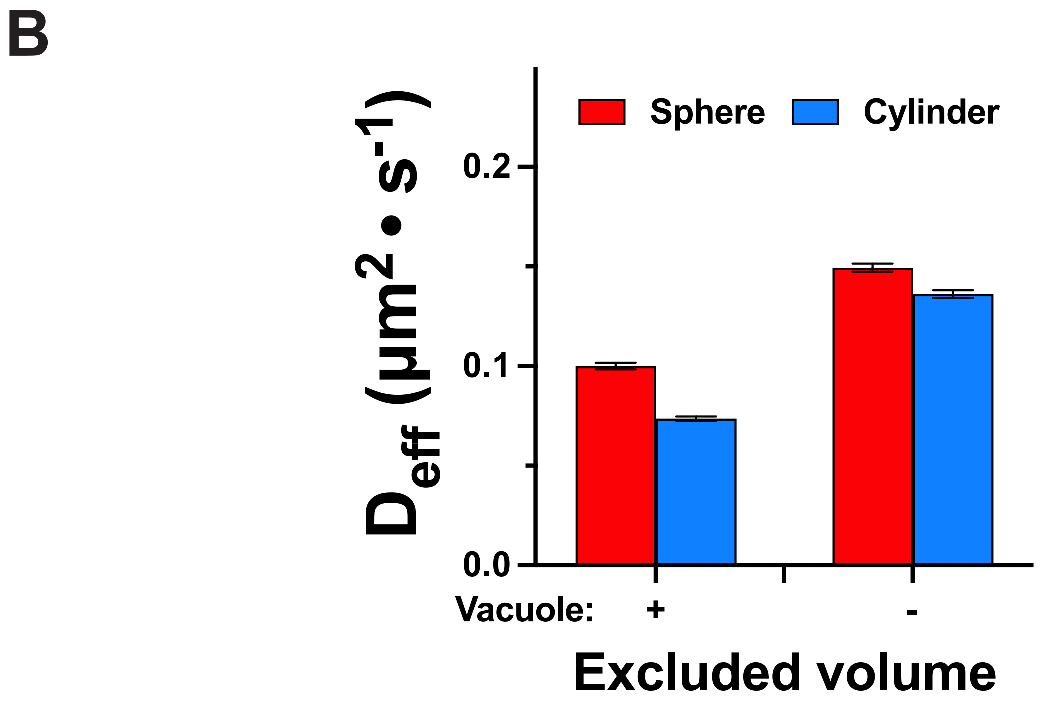
B
Sphere
Cylinder
$D_{eff}$ (µm$^2$ • s$^{-1}$)
0.2
0.1
0.0
Vacuole:
+
-
Excluded volume

Figure S5

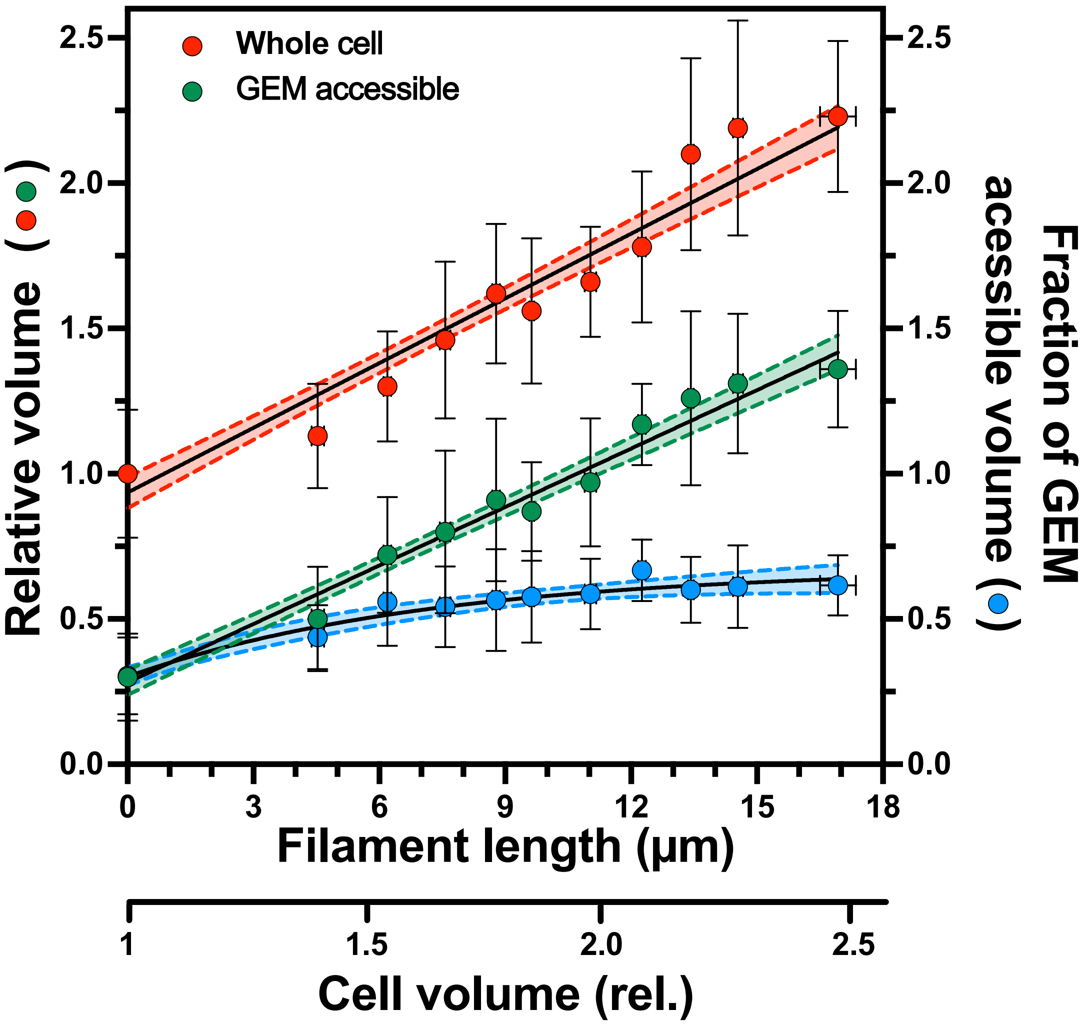

Whole cell
GEM accessible
Relative volume
Fraction of GEM acessible volume
Filament length (µm)
Cell volume (rel.)
0.0
0.5
1.0
1.5
2.0
2.5
0
3
6
9
12
15
18
1
1.5
2.0
2.5


Figure S6

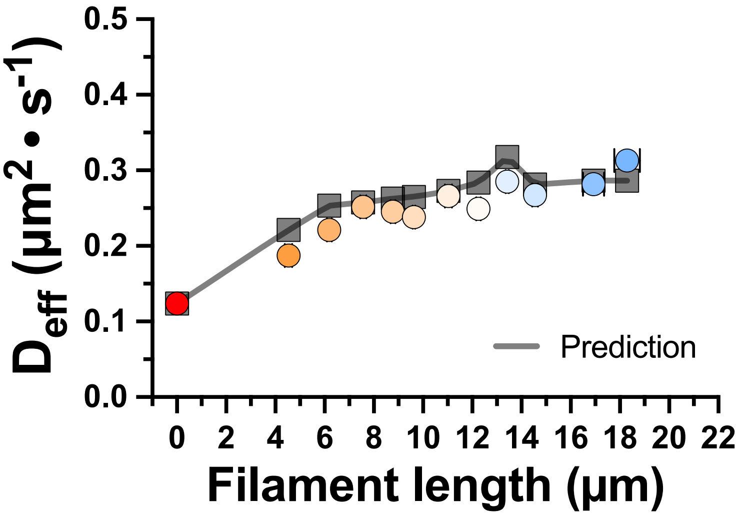
0.5
0.4
0.3
0.2
0.1
0.0
D_eff (µm² • s⁻¹)
Prediction
0 2 4 6 8 10 12 14 16 18 20 22
Filament length (µm)

**Figure S7**

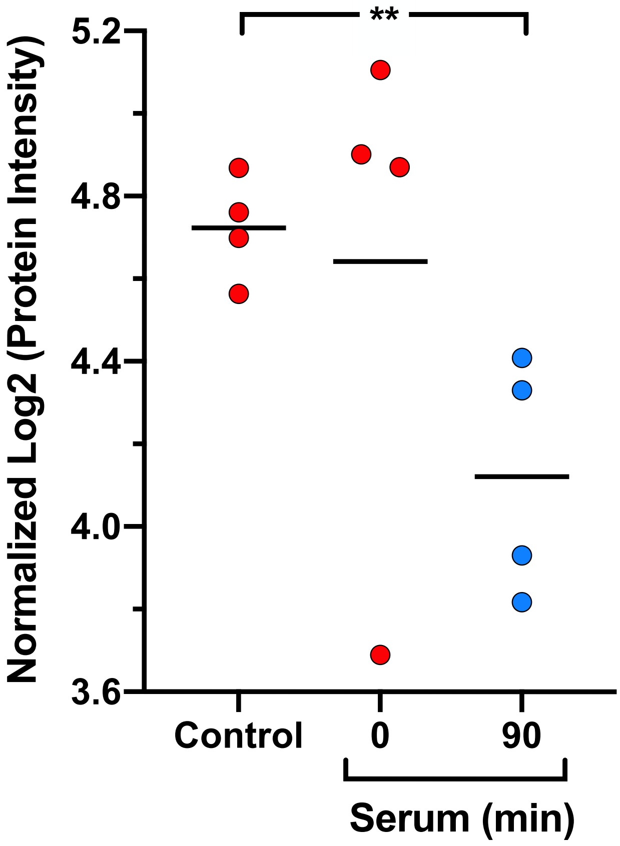
**
5.2
4.8
4.4
4.0
3.6
Normalized Log2 (Protein Intensity)
Control
0
90
Serum (min)

**Figure S8**

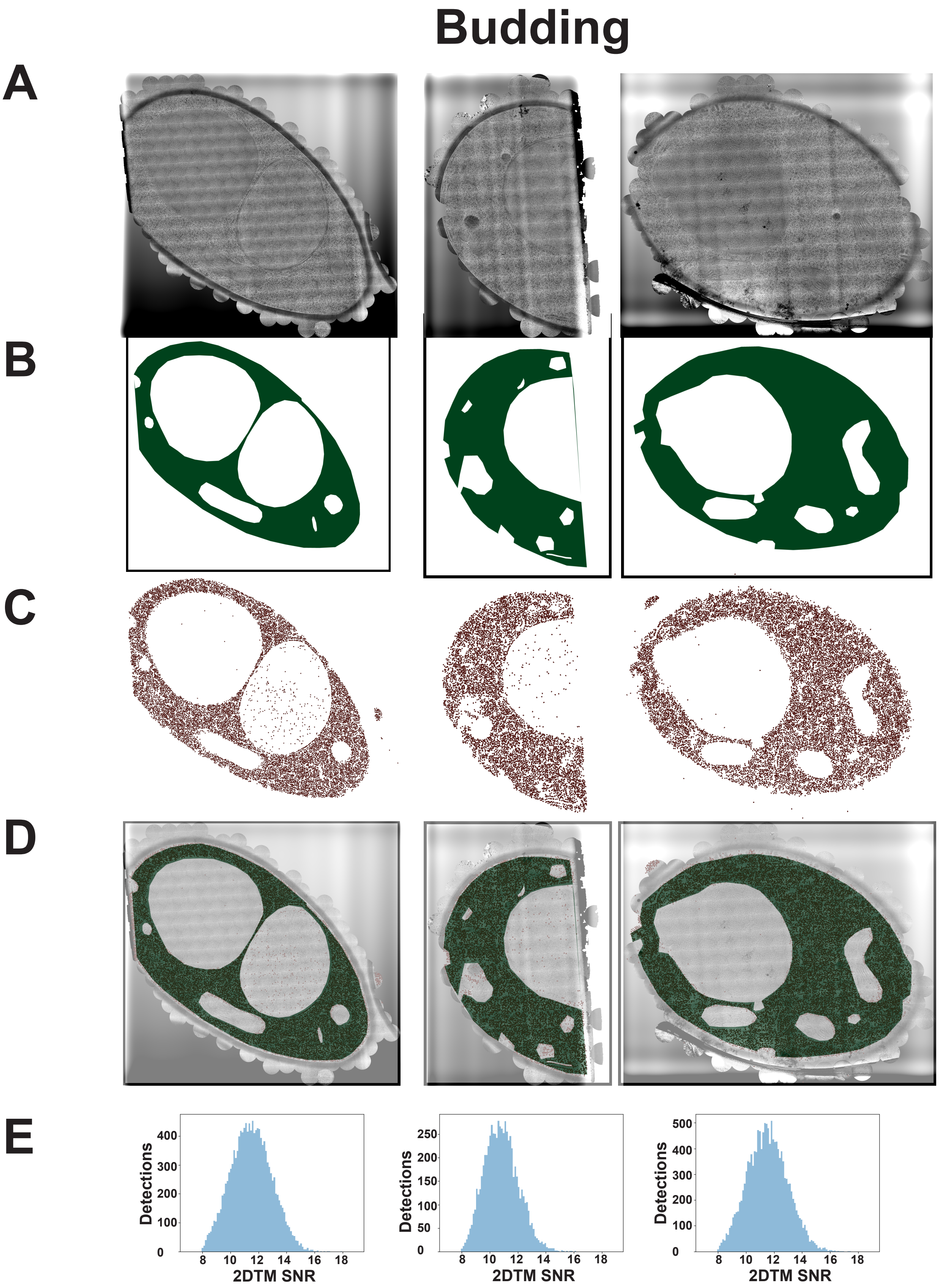

Budding
A
B
C
D
E
Detections
400
300
200
100
0
8 10 12 14 16 18
2DTM SNR
Detections
250
200
150
100
50
0
8 10 12 14 16 18
2DTM SNR
Detections
500
400
300
200
100
0
8 10 12 14 16 18
2DTM SNR


**Figure S9**

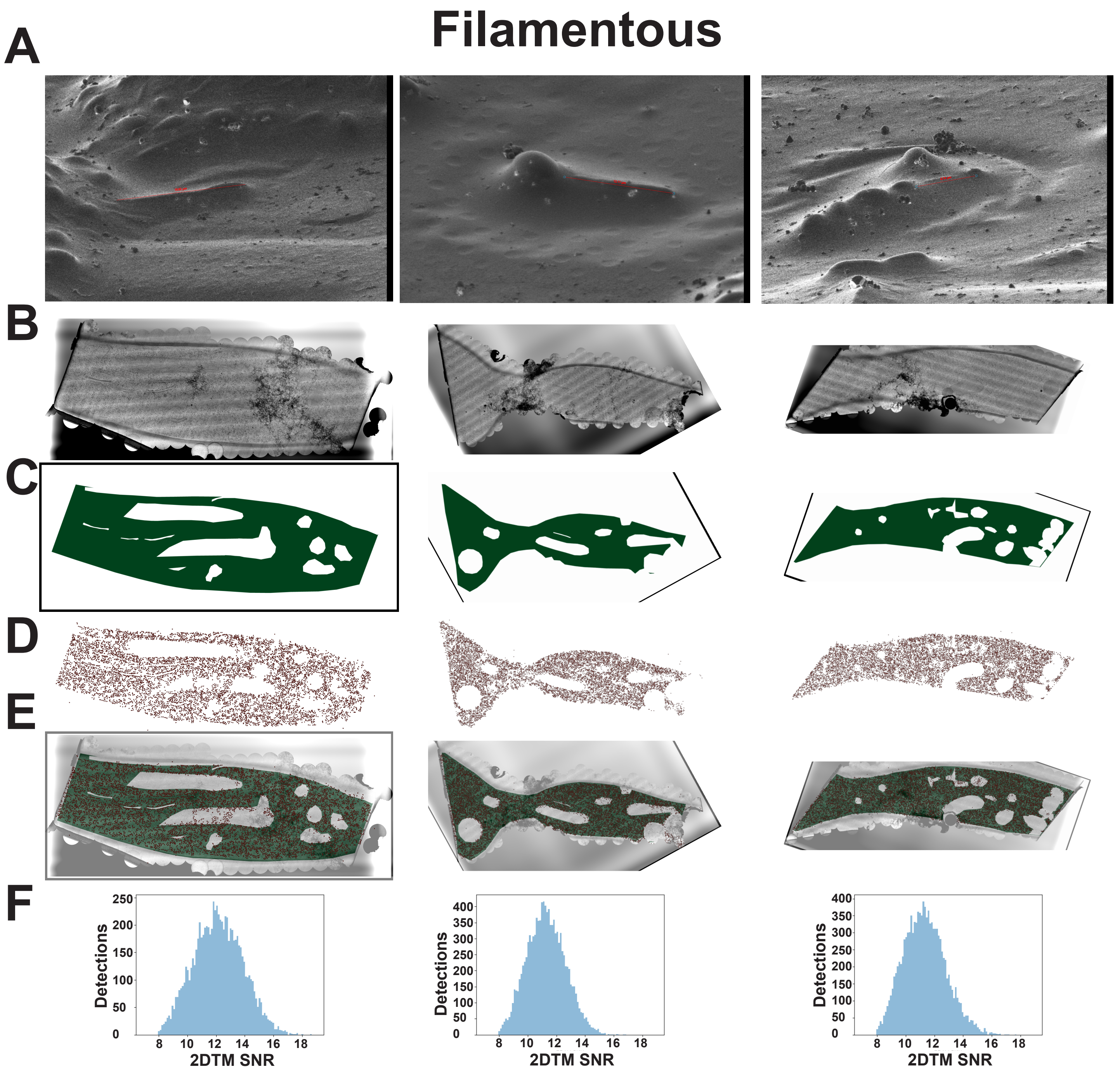
Filamentous
A
B
C
D
E
F
250
200
150
100
50
0
Detections
8 10 12 14 16 18
2DTM SNR
400
350
300
250
200
150
100
50
0
Detections
8 10 12 14 16 18
2DTM SNR
400
350
300
250
200
150
100
50
0
Detections
8 10 12 14 16 18
2DTM SNR

Figure S10

A

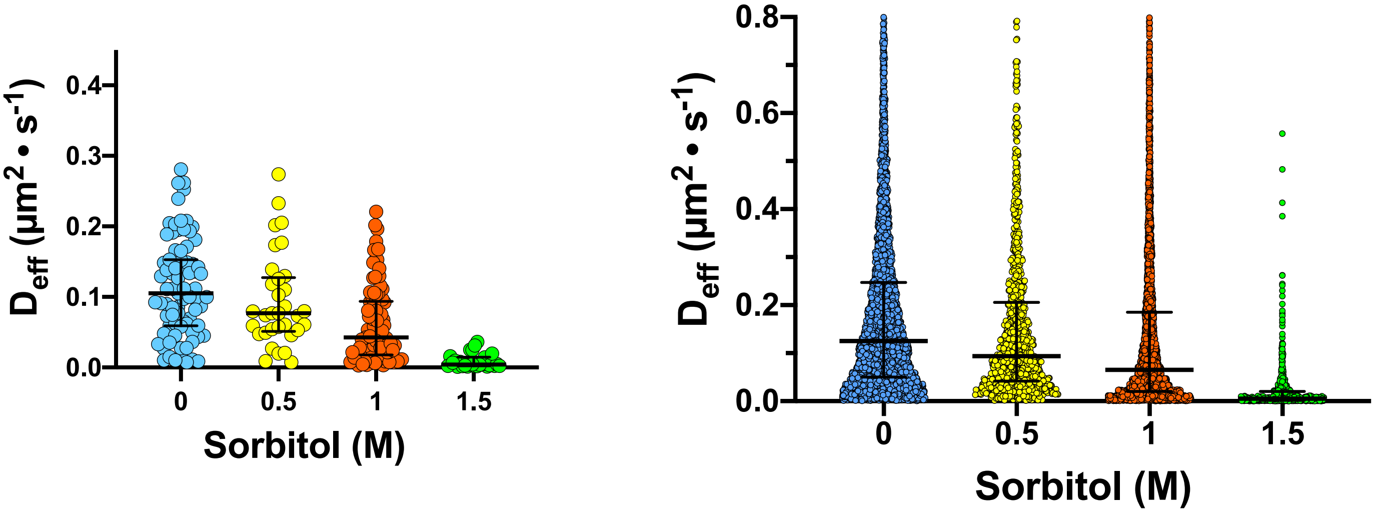


B

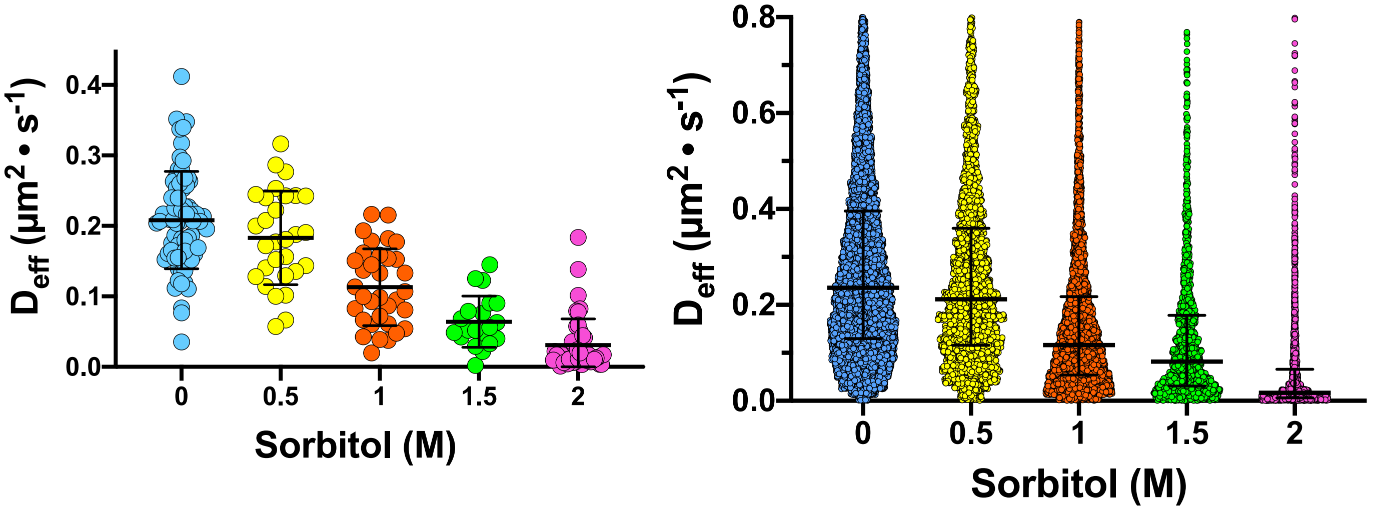


Figure S11

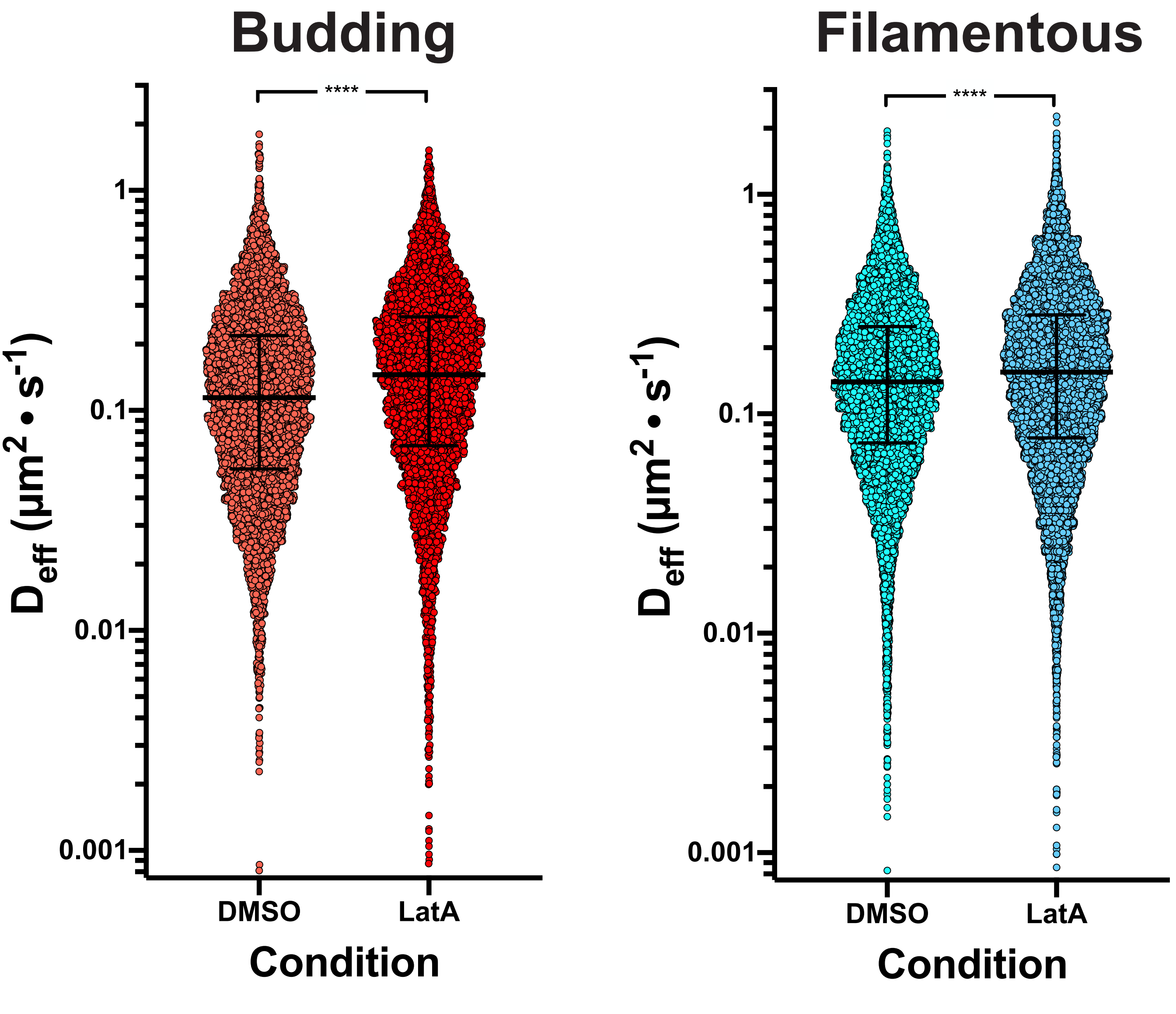

Budding
Filamentous
****
****
1
0.1
0.01
0.001
$D_{eff}$ (µm$^2$ • s$^{-1}$)
DMSO
LatA
Condition


**Figure S12**

A

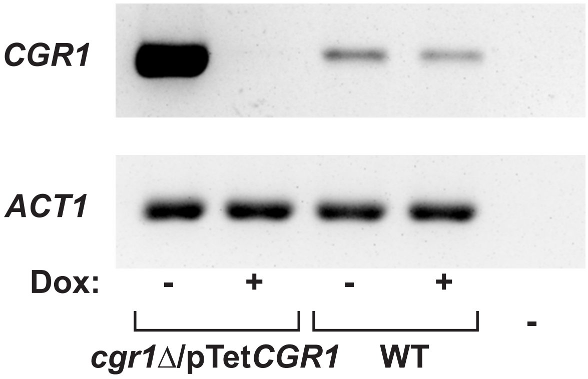


B

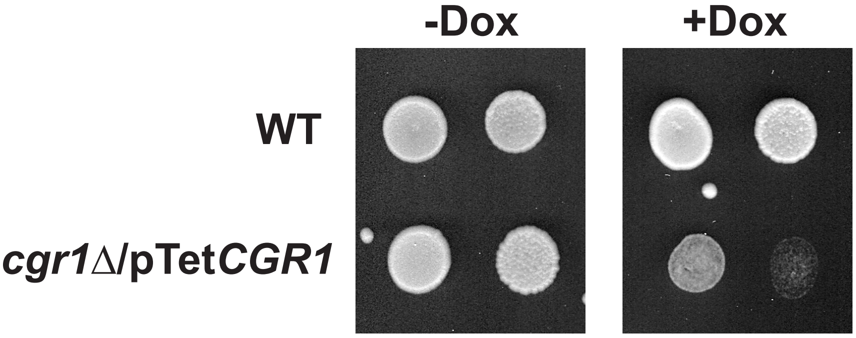


C

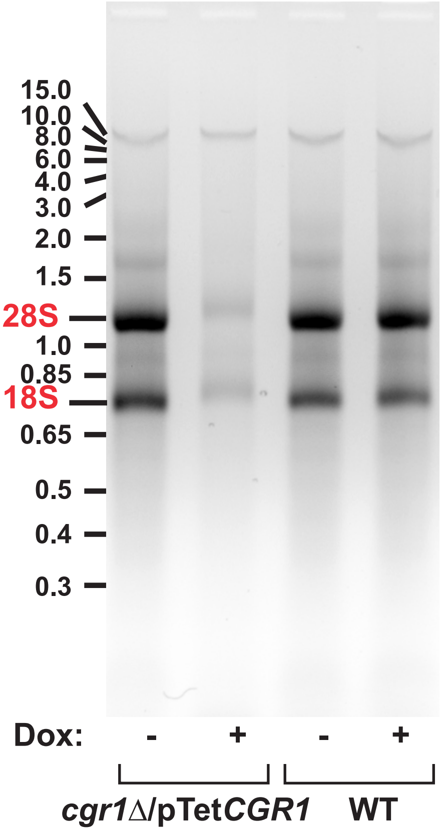


**Figure S13**
